# Fifty generations of amitosis: tracing asymmetric allele segregation in polyploid cells with single-cell DNA sequencing

**DOI:** 10.1101/2021.03.29.437473

**Authors:** Valerio Vitali, Rebecca Rothering, Francesco Catania

## Abstract

Amitosis is a widespread form of unbalanced nuclear division whose biomedical and evolutionary significance remain unclear. Traditionally, insights into the genetics of amitosis are acquired by assessing the rate of phenotypic assortment. The phenotypic diversification of heterozygous clones during successive cell divisions reveals the random segregation of alleles to daughter nuclei. Though powerful, this experimental approach relies on the availability of phenotypic markers. Here, we present an approach that overcomes the requirement for phenotypic assortment. Leveraging *Paramecium tetraurelia*, a unicellular eukaryote with nuclear dimorphism and a highly polyploid somatic nucleus, we use single-cell whole-genome sequencing to track the assortment of developmentally acquired somatic DNA variants. Accounting for genome representation biases, we measure the effect of amitosis on allele segregation across the first ∼50 amitotic divisions post self-fertilization and compare our empirical findings with theoretical predictions estimated via mathematical modeling. In line with our simulations, we show that amitosis in *P. tetraurelia* produces measurable but modest levels of *s*omatic assortment. In forgoing the requirement for phenotypic assortment and employing developmental, environmentally induced somatic variation as molecular markers, our work provides a new powerful approach to investigate the consequences of amitosis in polyploid cells.

## Introduction

The commonly held view that mitosis and meiosis are the universal forms of cell division is incomplete—some cells can also divide without the intervention of the nuclear spindle following direct nuclear fission, a process known as amitosis.

The existence of amitosis has been repeatedly called into question. Many of its early accounts (*e.g.* **(Child 1907)**) have been disproved **(Conklin 1917)**, its occurrence considered a rare exception **(Pfitzer 1980)**, an aberrant or degenerative process **(Flemming 1891)**, or a form of nuclear division strictly uncoupled from cell proliferation **(Macklin 1916)** and of uncertain functional significance. Since then, various forms of “true” amitosis have been documented across eukaryotes including insects **(Lucchetta and Ohlstein 2017; Nakahara 1917)**, plants **(Miller 1980)**, and more tentatively, vertebrates **(Kuhn, Therman, and Susman 1991; Yiquan and Binkung 1986)**. Most notably, in ciliates amitosis has evolved into the predominant means of somatic nuclear reproduction during cell proliferation **(Orias 1991)**.

In *Drosophila,* amitosis of polyploid cells in the intestinal epithelium may serve as a significant mechanism of de-differentiation associated with stem cell replenishment **(Lucchetta and Ohlstein 2017)**. This mechanism may also initiate cancer through the formation of aneuploid cells **(Lucchetta and Ohlstein 2017)**. In vertebrates, amitosis may occur in damaged or cancerous liver cells **(Yiquan and Binkung 1986)**, or in deciduous tissues with subpopulations of polyploid cells such as the trophoblast **(Kuhn et al. 1991)**. Polyploidy, achieved through endomitosis or endoreplication **(Fox and Duronio 2013; Zielke, Edgar, and DePamphilis 2013)**, may promote DNA-damage insensitivity through various mechanisms in plants, insects and bacteria, and serve as a virulence factor in pathogenic fungi **(Schoenfelder and Fox 2015)**. In addition, mitotic de-polyploidization of polyploid cells is associated with cell rejuvenation in cancer **(Erenpreisa et al. 2011)**, and, similar to amitosis, can readily generate populations of genetically heterogeneous cells (aneuploid cells) capable of rapid adaptive evolution (*e.g.* in response to xenobiotics or tissue damage **(Duncan et al. 2010, 2012)**). Despite the widespread phylogenetic distribution of amitosis, its potential role in stem cell differentiation, and cancer onset and progression, this form of unbalanced nuclear division is severely understudied.

Ciliates offer a powerful system for gaining insights into the process of amitosis. Ciliated protozoans such as *Paramecium tetraurelia* (henceforth *Paramecium*) are characterized by two functionally specialized nuclei with distinct nuclear architectures **(Lyn 2010)**. The small diploid germline nucleus, the micronucleus, is transcriptionally silent during asexual division and harbors the germline genome. In contrast, the larger somatic nucleus—the macronucleus—is expressed during vegetative growth. Its expression governs cell physiology and behavior **(Beale and Preer Jr. 2008a)**. In *Paramecium*, the somatic genome is highly polyploid. This high-level ploidy is achieved during the biogenesis of the macronucleus through an endoreplication process, in which a copy of the diploid germline genome is used as a template for amplification (from 2*n* to ∼860*n* **(Allen and Gibson 1972; Woodard, Gelber, and Swift 1961)**).

During the vegetative life of *Paramecium*, the diploid micronuclei divide mitotically, whereas the polyploid somatic nucleus divides amitotically—it elongates and eventually separates into two daughter macronuclei. Upon amitosis allele segregation is subject to random fluctuations. It is not entirely clear how cells can avoid severe aneuploid imbalances over prolonged vegetative division **(Preer and Preer 1979)**. This is especially true for the ciliate *Tetrahymena*, which has a much lower ploidy than *Paramecium* (∼45*n* **(Doerder, Deak, and Lief 1992; Eisen et al. 2006; Hamilton et al. 2016; Orias and Flacks 1975)**). Although not necessarily sufficient to maintain constant ploidy levels across the genome, there is evidence that in *Paramecium* the total macronuclear DNA content is tightly regulated across divisions **(Berger and Schmidt 1978)**. This hints at the existence of a compensatory “replicative control” mechanism that may occur at the individual chromosome level **(Beale and Preer Jr. 2008b; Preer and Preer 1979)**. Such a mechanism would prevent aneuploid imbalance (deviations from the original ploidy) or even complete chromosomal loss (so called *nullisomics*, where both alleles are lost). Alternatively, as suggested by a more recent study on the ciliate *Chilodonella uncinata*, balancing selection may be sufficient to maintain a stable ploidy during asexual reproduction **(Spring, Pham, and Zufall 2013)**.

Due to the random assortment of genetic elements during amitosis of the macronucleus (henceforth *somatic assortment*), a biallelic locus eventually becomes fully homozygous for either of the alternative alleles. The rate at which this loss of heterozygosity occurs is primarily determined by the number (ploidy) and nature of the segregating units and the input ratio, *i.e.*, the relative proportion of the two somatic alleles at the beginning of the clonal cycle **(Bell 2009; Doerder et al. 1992; Merriam and Bruns 1988)**. Because ciliates’ macronuclei determine the cell phenotype, *somatic assortment* at heterozygous loci may give rise to *phenotypic assortment*—heterozygous clones eventually segregate into homozygous sub-clones stably expressing one of the two parental alleles **(Doerder et al. 1992; Merriam and Bruns 1988; Nanney and Preparata 1979; Orias and Flacks 1975)**. *Phenotypic assortment* has been the primary tool for investigating *somatic assortment* and has greatly helped understand the nature of amitosis in ciliates such as *Tetrahymena* **(Doerder et al. 1992)**. However, a simple and direct approach that helps illuminate the process of amitosis that does not rely on phenotypic traits is currently lacking. Such an approach would conveniently allow researchers to investigate amitosis even in the absence of genetic markers that encode easily observable traits.

Recent findings concerning the process of soma development in *Paramecium* open a new perspective on how amitosis can be studied. Like other ciliates, the polyploid somatic genome of *Paramecium* is an extensively processed version of the germline genome, largely deprived of a considerable portion of DNA via a developmental process called Programmed DNA Elimination (PDE). In addition to removing transposons and other repetitive DNA elements, PDE removes tens of thousands of intervening, typically short (<150bp) and AT-rich germline DNA elements termed Internal Eliminated Sequences (IESs) **(Arnaiz et al. 2012; Beale and Preer Jr. 2008a; Duharcourt and Betermier 2014; Guérin et al. 2017)**. Although IESs are, for the most part, perfectly removed from the newly developed somatic genome, some are incompletely excised—in the order of a few hundreds at standard cultivation conditions **(Vitali, Hagen, and Catania 2019)**. These retained elements, which we termed somatic IESs, interrupt a variable fraction (henceforth retention levels) of the total number of macronuclear DNA copies **(Arnaiz et al. 2012; Catania et al. 2013; Duret et al. 2008; Hagen, Vitali, and Catania 2020; Vitali et al. 2019)**. The retention levels of somatic IESs provide a measurable molecular marker to assess the random assortment of segregating alleles in *Paramecium*. More explicitly, by recording the retention levels of somatic IESs across subsequent amitotic cycles (*i.e.*, asexual generations), it should be possible to directly test the extent to which amitosis impacts the segregation of somatic alleles.

Single-cell sequencing technology (scDNA-seq) is a potentially powerful approach to test this idea. The reliable detection of amitosis-associated changes in allele frequencies necessitates deep and comprehensive genome coverage as well as sensitivity and faithfulness. After individual cell isolation, scDNA-seq protocols invariably involve a step of extensive whole genome amplification (WGA) followed by library construction and next generation sequencing of the amplification products. Depending on the specific amplification technology and application, the WGA step can produce a satisfactory representation of the target genome **(Huang et al. 2015; Pinard et al. 2006)**. However, WGA may also result in amplification artifacts, such as overrepresentation of large templates **(Maurer-Alcalá, Knight, and Katz 2018; Sabina and Leamon 2015)**, reduced genome coverage **(Börgstrom et al. 2017)**, misrepresentation of copy number variants under certain conditions **(Van Der Plaetsen et al. 2017)** (but see (**(Deleye et al. 2017)**)), poor scaffold assembly **(de Bourcy et al. 2014)**, and allele dropout **(Luquette et al. 2019)**. Current commercially available kits for non-PCR based single-cell WGA minimize amplification artifacts through a highly optimized isothermal Multiple Displacement Amplification (MDA) reaction **(Meier et al. n.d.; Pinard et al. 2006)**. Although MDA-based WGA is far more resilient to genome representation biases compared to thermocycling methods **(Lasken and Egholm 2003)**, it may preferentially amplify GC-rich regions **(Sabina and Leamon 2015)** and lead to an underrepresentation of AT-rich regions (*e.g. Paramecium*’s IESs). This “selection bias” is anticipated to reach concerning levels in organisms whose genome composition lies at the low end of the GC-spectrum, such as fungi, amoebas, apicomplexans, and ciliates **(Videvall 2018)**. Potential caveats aside, scDNA-seq could be a powerful tool to trace stochastic evolution in amitotically-dividing cells.

Here we leverage the advantages, and probe the limits of, scDNA-seq to investigate the assortment of somatic IESs in individual *Paramecium* cells across successive amitotic divisions. In addition, we develop a freely available software, which simulates the random segregation of genetic elements across amitotic divisions **(Vitali, Hagen, and Catania 2021)**, and determine the theoretical rate of somatic assortment in *Paramecium*. By comparing empirical data with simulation-based predictions, we find that amitosis-associated changes in allele frequencies in *Paramecium* deviate modestly from what is expected under random assortment. Collectively, we show that single-cell whole genome sequencing and dedicated bioinformatic analyses allow accurate tracing of amitotic allele segregation in proliferating polyploid cells.

## Results

### Single-cell DNA sequencing of the *Paramecium* somatic genome

scDNA-seq should facilitate direct measurements of the fraction of segregating alleles in individual somatic nuclei of asexually reproducing cells. Among the available scDNA-seq options, the Multiple Displacement Amplification (MDA)-based methods generate the highest amplification yield and most complete genome coverage while introducing minimal bias relative to other amplification methods **(Lasken and Egholm 2003; Meier et al. n.d.; Pinard et al. 2006)**. However, how extensively genomes or genomic regions with particularly low GC content—such as the somatic DNA of *Paramecium* (28% on average)—prove refractory to MDA has not yet been ascertained to our knowledge.

To assess the quality of the scDNA-seq data in terms of somatic genome representation and coverage, we compared a total of 11 scDNA samples to a mass culture sample (mcDNA) obtained from the parental population used to set up the single-cell experiment. Additionally, we included a computer-generated DNA-seq sample (artificial DNA, aDNA) produced from *P. tetraurelia*’s reference (somatic) genome to serve as bias-free reference.

All scDNA-seq samples examined show a moderate underrepresentation of AT-rich sequences (**Figure 1A**). In contrast, both the mcDNA-seq and aDNA-seq show virtually homogeneous coverage across the whole range of GC-content found in the *Paramecium* genome. Furthermore, sequencing depth in the scDNA-seq samples increases with the distance from scaffold ends (**Figure 1B**). This observation suggests that there is a substantial reduction of amplification efficiency of the MDA reaction at the chromosome termini. A quantitative analysis of genome representation confirms that scDNA samples suffer from moderate to intermediate *GC Bias*, *i.e.*, the underrepresentation of AT-rich regions, and severe *Terminal Bias*, *i.e.*, the underrepresentation of chromosome termini (**Table 1**).

**Figure 1.**
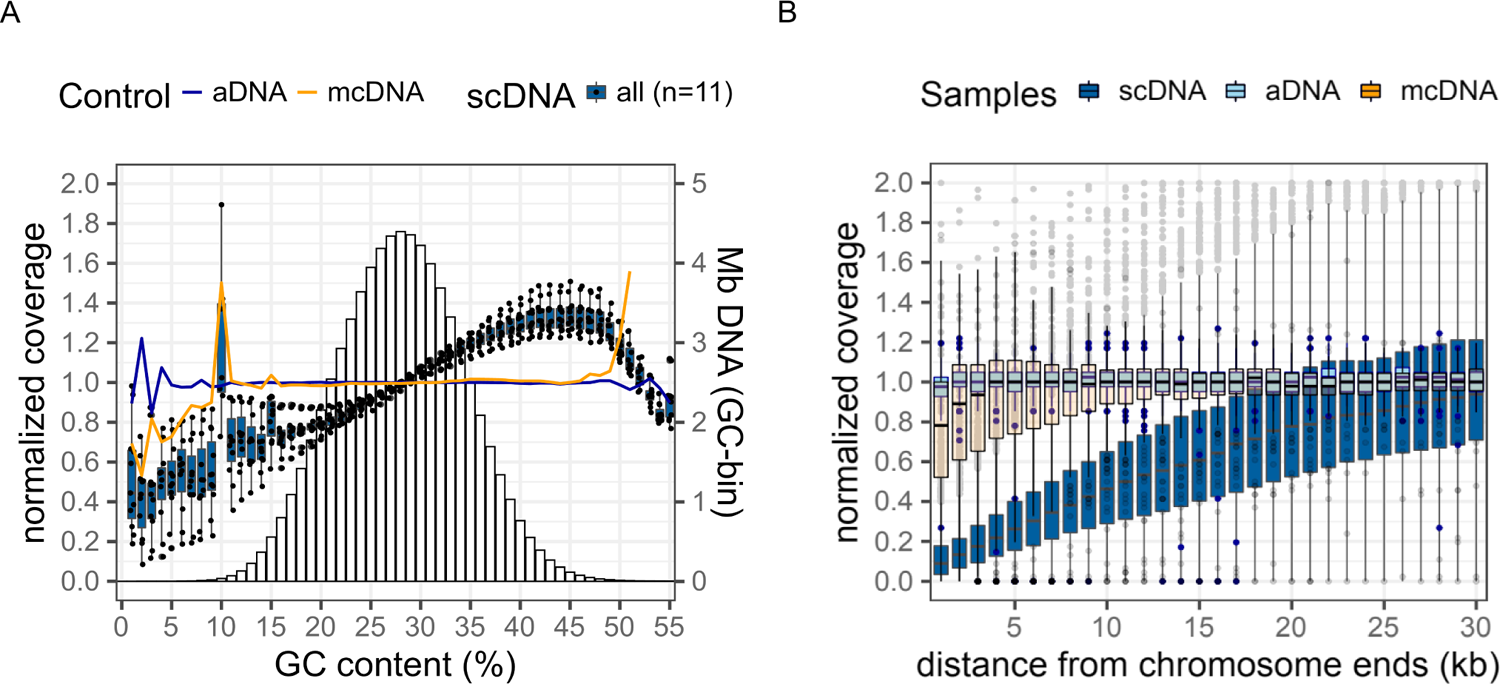
Amplification biases of MDA-based single-cell DNA seq. A) Positive GC bias. Change in normalized base coverage with GC content (%). Normalized coverage = n° reads / base / mean coverage. Bar chart in the background shows the amount of DNA for each GC bin (Megabases, Mb, secondary axis). **B) Terminal representation bias.** Change in normalized base coverage with distance from chromosome termini (kilobases, kb). The degree of GC bias and underrepresentation of scaffold ends are shown for 11 single-cell sequencing samples (scDNA), their parental mass culture sample (mcDNA) and one artificially generated sample (aDNA). MDA, Multiple Displacement Amplification. scDNA, single-cell DNA sequencing. mcDNA, mass culture DNA sequencing. aDNA, artificial DNA sequencing.

**Table 1.** Quantitative analysis of genome representation. GC Bias. Linear regression of normalized coverage on GC content. GC bias estimates are expressed as change of normalized coverage every 10% change in GC content. Normalized coverage is shown for DNA with GC content one standard deviation (sd) above (∼22%) and below (∼34%) the mean (28% GC). Perc., percentile. *b*, regression coefficient. **Terminal Bias.** Linear regression of normalized coverage on distance from chromosome ends (every 10kb). True relationship is parabolic. Normalized coverage is estimated for regions that are 1 and 30 kb away from either chromosome ends. Terminal Bias was calculated on the 115-telomere-capped chromosomes of *P. tetraurelia*. aDNA-seq, artificially-generated DNA sequencing. mcDNA-seq, mass culture DNA sequencing. scDNA-seq, single-cell DNA sequencing. Mean ± sd of the mean is shown for 11 scDNA-seq samples.

| Sample | GC Bias |  |  | Terminal Bias |  |  |
| --- | --- | --- | --- | --- | --- | --- |
|  | *b* | Coverage |  | *b* | Coverage |  |
|  |  | 16th perc. (22%<br>GC) | 84th perc. (34%<br>GC) |  | 1kb away | 30kb away |
| **aDNA** | 0.001 | 0.999 | 1.000 | 0.006 | 0.981 | 1.007 |
| **mcDNA** | -0.009 | 0.985 | 1.010 | 0.059 | 0.830 | 1.000 |
| **scDNA** | 0.163 ± 0.059 | 0.811 ± 0.043 | 1.160 ± 0.036 | 0.321 ± 0.025 | 0.140 ± 0.024 | 1.037 ± 0.029 |

### Detection of AT-rich germline sequences in the *Paramecium* somatic genome

Somatic IESs may be viewed as AT-rich insertions that occur naturally in *Paramecium* following somatic genome development **(Arnaiz et al. 2012; Catania et al. 2013; Duret et al. 2008; Hagen et al. 2020; Vitali et al. 2019)**. Detection of these somatic IESs requires pervasive and deep genome coverage as mutant alleles (IES^+^) are scattered across the genome and can be retained in a variable fraction of the polyploid somatic nucleus, coexisting with their wild-type alleles (IES^−^). To determine whether the uncovered biases of scDNA-seq (**Figure 1** and **Table 1**) limit our ability to detect somatic IESs, we compared the somatic genomes obtained from mass culture and single cells. It is worth noting that, unlike scDNA, conventional mcDNA (bulk) sequencing does not capture the genetic heterogeneity of single cells, and for a given locus provides a population-average estimate of the fraction of target somatic alleles.

Relative to the reference mcDNA, scDNA samples with a similar number of mapped reads (scDNA_1x) exhibit higher levels of IES dropout (*i.e.*, poor or no coverage of IES-flanking macronuclear regions) due to uneven genome representation (**Table 2**). However, this effect is ameliorated by increased sequencing depth (**Figure 2A**), and scDNA samples with approximately double the amount of mapped reads (scDNA_2x) show IES dropout levels comparable to those of the reference mcDNA (**Table 2**). When we account for the level of total IES dropout, the number of somatic IESs inferred for the scDNA samples approaches the number of somatic IESs detected in the reference mcDNA sample (**Figure 2B**). Last, we tested whether IES retention levels, as measured through the IES Retention Score (IRS, see Methods), are underestimated in scDNA samples as compared to the mcDNA sample. Despite the elevated AT content of IESs and the detected GC bias associated with single-cell DNA sequencing, we don’t find evidence for preferential dropout of the mutant allele (IES^+^) (**Additional File 1: Figure S1**). Overall, we show that MDA-based scDNA-seq, when applied to an AT-rich genome such as that of *Paramecium* can yield comprehensive genome coverage as long as sequencing depth is sufficiently large, ideally >2 fold compared to mass culture sequencing (**Additional File 1: Table S1**).

**Figure 2.**
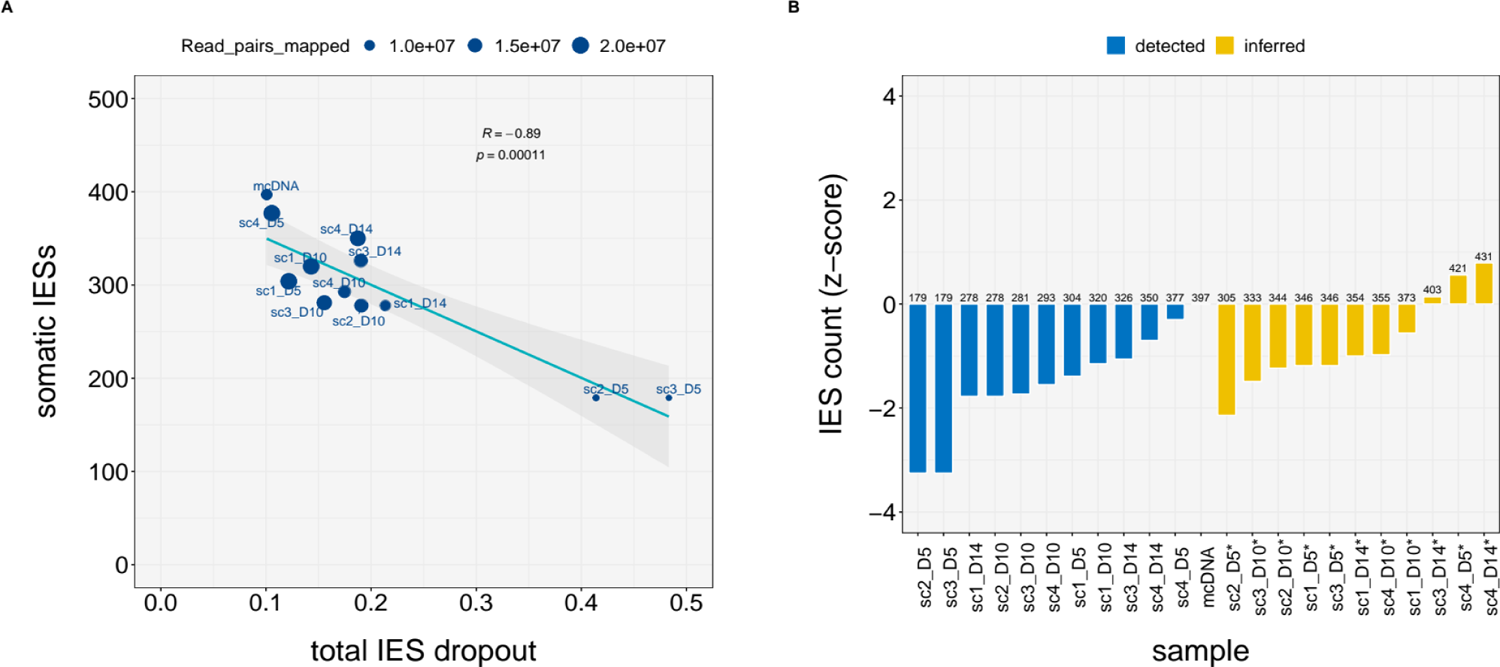
IES dropout due to uneven genome representation in scDNA samples. A) Number of detected somatic IESs as a function of coverage. Number of somatic mutations detected as a function of Total IES dropout (“invisible” IES loci) and number of read pairs mapped (dot size). Somatic IESs ∼ Total IES dropout, r = 0.882, *P* < 0.01. **B) Count of somatic IESs before and after correction.** Somatic IES counts before and after correcting for Total IES dropout. Deviation is relative to the count obtained for bulk DNA-seq (mcDNA; z-score = 0). Correction, count / (1 − Total IES dropout). Deviation from mcDNA count, IES count *z-score* = (IES_count *−* ref_value) / sd. Counts and corrected counts are indicated above bars. Sample names for corrected counts are labeled with a star sign.

**Table 2.** Quantitative analysis of IES dropout. Total dropout. Fraction of all known IES loci (*n*=44,928) with read coverage equal to or lower than 20. **Terminal dropout.** Fraction of all known IES loci located within 30 kb from either scaffold ends (*n*=9,986) with a read coverage equal to or lower than 20. **GC dropout.** IES dropout unexplained by either terminal or residual dropout is assumed to results from the positive GC Bias. **Residual dropout.** IES dropout unrelated to amplification biases found in the mcDNA sample. Mapped pairs, total number of read pairs mapped (in millions). mcDNA, mass culture DNA sequencing. scDNA, single-cell DNA sequencing. scDNA_1x, scDNA samples with approximately the same number of mapped reads compared to the mcDNA sample (5*10^6 < *n*° mapped reads < 15*10^6, *n*=6). scDNA_2x, scDNA samples with approximately twice as many mapped reads compared to the mcDNA sample (*n*° mapped reads > 19*10^6, *n*=4). Mapped, mapped read pairs (Millions).

| Sample | Mapped (M) | IES dropout |  |  |  |
| --- | --- | --- | --- | --- | --- |
|  |  | Total | Terminal | GC | Residual |
| mcDNA | 10.92 | 0.10 | 0.05 | 0.00 | 0.06 |
| scDNA_1x | $11.29 \pm 3.62$ | $0.28 \pm 0.14$ | $0.11 \pm 0.02$ | $0.11 \pm 0.09$ | $0.06 \pm 0.02$ |
| scDNA_2x | $19.67 \pm 0.51$ | $0.12 \pm 0.02$ | $0.08 \pm 0.01$ | $0.02 \pm 0.01$ | $0.03 \pm 0.001$ |

### IES retention levels across the first ∼50 amitotic divisions post self-fertilization

Having assessed the quality of the scDNA-seq data and learned how to mitigate the impact of scDNA-seq biases, we examined progressively aging *Paramecium* lines and estimated IES retention scores (IRSs) for cells collected on Day 5 (4 replicates), Day 10 (4 replicates) and Day 14 (3 replicates). These cells had undergone, respectively, ∼17, ∼35 and ∼49 divisions after the last self-fertilization. We focused on a set of highly covered IES loci (*N*=75) for which we could accurately estimate the corresponding retention levels. We asked: how do the empirical IRS values change over time?

We compared the changes in the standard deviation for the empirical IRS values (observed SDIRS), and their ratios (SDRIRS) across time points (**Additional File 1: Figure S2**) We find a significant up-shift in the SDIRS distribution over time (**Additional File 1: Figure S2A**) when comparing the two points furthest apart in the time course (Wilcoxon signed rank test, D14 vs. D5, *P*= 0.037, effect size r = 0.282 (small), *N* = 75). When considering the standard deviation ratios (SDRIRS) computed pairwise between time points, the difference is only slightly above the significance threshold (Wilcoxon signed rank test, D14 / D5 vs. D10 / D5, *P* = 0.062, effect size r = 0.199 (small), *N* = 60), although the density plots show a clear up-shift in the distribution over time (median D14 / D5 SDR_IRS_ = 1.25) (**Additional File 1: Figure S2B**). We also report the summary statistics for the observed and predicted IRS standard deviations (**Additional File 1: Table S2**). Taken together, our empirical findings suggest a slight increase in variation of IES retention levels across amitotic cell divisions.

### Simulation of somatic assortment

Multiple models of macronuclear architecture in ciliates have been proposed to account for observed rates of phenotypic assortment, the relative difference in DNA content between micro- and macronuclei, the absence of visible mitosis, and the avoidance of aneuploid imbalance. However, most if not all models proposed so far suffer from some sort of limitations **(Bell 2009; Nanney and Preparata 1979; Preer and Preer 1979)**. Three fundamental macronuclear configuration models (alongside their implications) are described in **Figure 3**. Briefly, the *chromosomal model* assumes that individual somatic chromosomes segregate independently from each other at cell division (**Figure 3A**), whereas the *diploid model* posits that homologous chromosomes (or set of chromosomes) are bundled into diploid sub-units (**Figure 3B**). Finally, the *whole-genome haploid sub-unit model* (hereafter the *haploid model*) hypothesizes that full sets of chromosomes from either one of the parental haplotypes are held together into larger segregating sub-units (**Figure 3C**). Provided that *Paramecium* avoids aneuploid imbalance at all loci, regardless of the mechanism **(Beale and Preer Jr. 2008a; Bell 2009; Berger and Schmidt 1978; Preer and Preer 1979)**, both the *chromosomal* and *haploid* models predict that for a somatic locus that retains an IES after Programmed DNA Elimination, the fraction of IES^+^ copies (mutant allele) will tend towards either 1 (IES Retention Score [IRS] = 1, only IES^+^ copies) or 0 (IRS = 0, only IES^−^ copies) as cells continue to divide asexually. But how rapidly would this loss of heterozygosity occur? To the best of our knowledge, while a thorough quantitative exploration of somatic assortment for *Paramecium* was published >40 years ago **(Preer 1976)**, direct evidence that the individual segregating subunits are somatic chromosomes (germline chromosome fragments) is currently lacking **(Nyberg 1976; Preer 1976)**.

**Figure 3.**
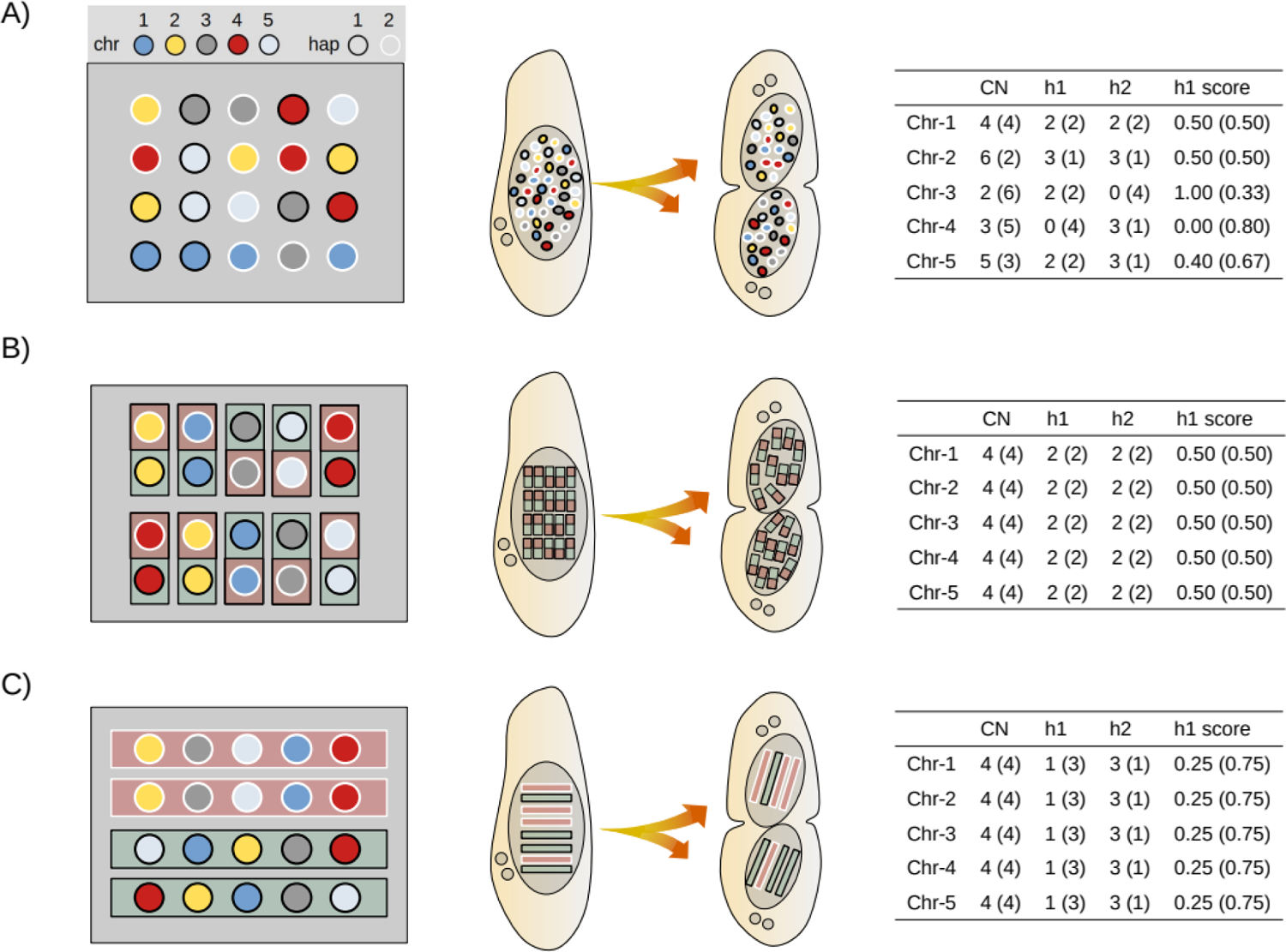
Models of macronuclear architecture in ciliates. Models for a hypothetical tetraploid cell with 5 somatic chromosome types (Chr) generated by conjugation (ex-conjugant). **Left.** Configuration of macronuclear sub-units in G1 (prior to DNA replication). **Center.** Random segregation of sub-units during amitotic division. **Right.** Copy number variation of individual chromosomes and their haplotypes after a single cell division. **A)** *Chromosomal model*. Individual somatic chromosomes segregate freely. N = 2 * Chr * k, where N is the total number of segregating sub-units at cell division and k is the ploidy level. **B)** *Diploid subunit model*. Homologous chromosomes are bundled up into diploid sub-units. N = Chr * k. **C)** *Whole-genome haploid subunit model*. Full sets of chromosomes are bundled into single haploid sub-units. Each sub-unit contains a full complement of chromosomal variants from either one of the parental haplotypes (but not both). N = 2 * k. CN, Copy Number. h1, CN of haplotype 1. h2, CN of haplotype 2. h1 score, nuclear prevalence of haplotype 1. h1 = h1 / (h1 + h2). Before cell division, CN = k and h1 = h2 = 0.50 for each chromosome type. Each daughter cell receives exactly half of the sub-units (N / 2) at cell division (number of sub-units in G1). All chromosomes are depicted as heterozygous for illustration purpose only.

We first used mathematical modeling to determine how the fraction of mutant alleles (IES^+^ copies) in the somatic nuclei is expected to change across successive amitotic divisions at individual IES loci. We simulated somatic assortment using the *haploid* and *chromosomal* models published by John Preer Jr. in 1976 **(Preer 1976)**. We used similar parameters, except for the number of somatic chromosomes, which was then assumed to be ∼43 **(Preer 1976)**, that we now know to be much larger due to chromosome fragmentation during DNA elimination. We set this parameter to 115, as there are 115 telomere-capped chromosomes in *Paramecium*’s genome annotation (but its number could be much larger, as 697 scaffolds larger than 2 kb were assembled) **(Aury et al. 2006)**. Our predicted values strongly correspond with those published by Preer, with only a slight discrepancy when running the simulation with the *chromosomal model* (**Additional File 1: Table S3**). To further validate our mathematical predictions we modeled somatic assortment for mass culture and daily re-isolation through bioinformatic simulations (see Methods). Mathematical and bioinformatics modeling have identical outcome (**Additional File 1: Figure S3**). The allele frequency variance for a small number of daily re-isolated lines follows a stochastic trend across generations. However, the average run for a large number of isolation cultures converges on the mathematical / mass culture predictions (**Additional File 1: Figure S3**). We provide new equations to calculate the standard deviation of allele frequency distributions (*e.g.* retention levels) and the rate of somatic assortment (*dσ/dt*) as a function of the number of asexual divisions and starting retention levels (Methods, equation (5-6)).

As expected, the simulation predicts an increase in variability of the copy number distribution of alleles (*e.g.* IES+ / IES^-^ copies) across generations (**Figure 4A**). The rate of somatic assortment is predicted to peak at an input ratio of 0.5 (starting retention level, IRS0 = 0.5), and decrease symmetrically around this value (**Figure 4B**). But how long would it take for the cells to experience a substantial loss of heterozygosity as a consequence of the random segregation of alleles at cell division? The simulation predicts that with a starting retention level (IRS0) of 0.5, after 200 asexual divisions (which corresponds roughly to a full clonal cycle of *Paramecium*), all cells would still be in the heterozygous state (IES^+^ and IES^−^ copies co-existing in the same nucleus) (**Figure 4A, red line and Figure 4C, inset**). In fact, somatic assortment of IES^+^ and IES^−^ alleles would only lead to a substantial loss of heterozygosity (e.g. *H* << 0.5) after thousands of asexual generations (**Figure 4C**). Furthermore, even when starting from IRS0 = 0.1 (or 0.9) the probability that an IES locus becomes fully homozygous after 200 divisions is smaller than 0.20 (**Figure 4C, inset, and Figure 4D**). In sum, our simulations predict that IES retention levels remain fairly stable during asexual division.

**Figure 4.**
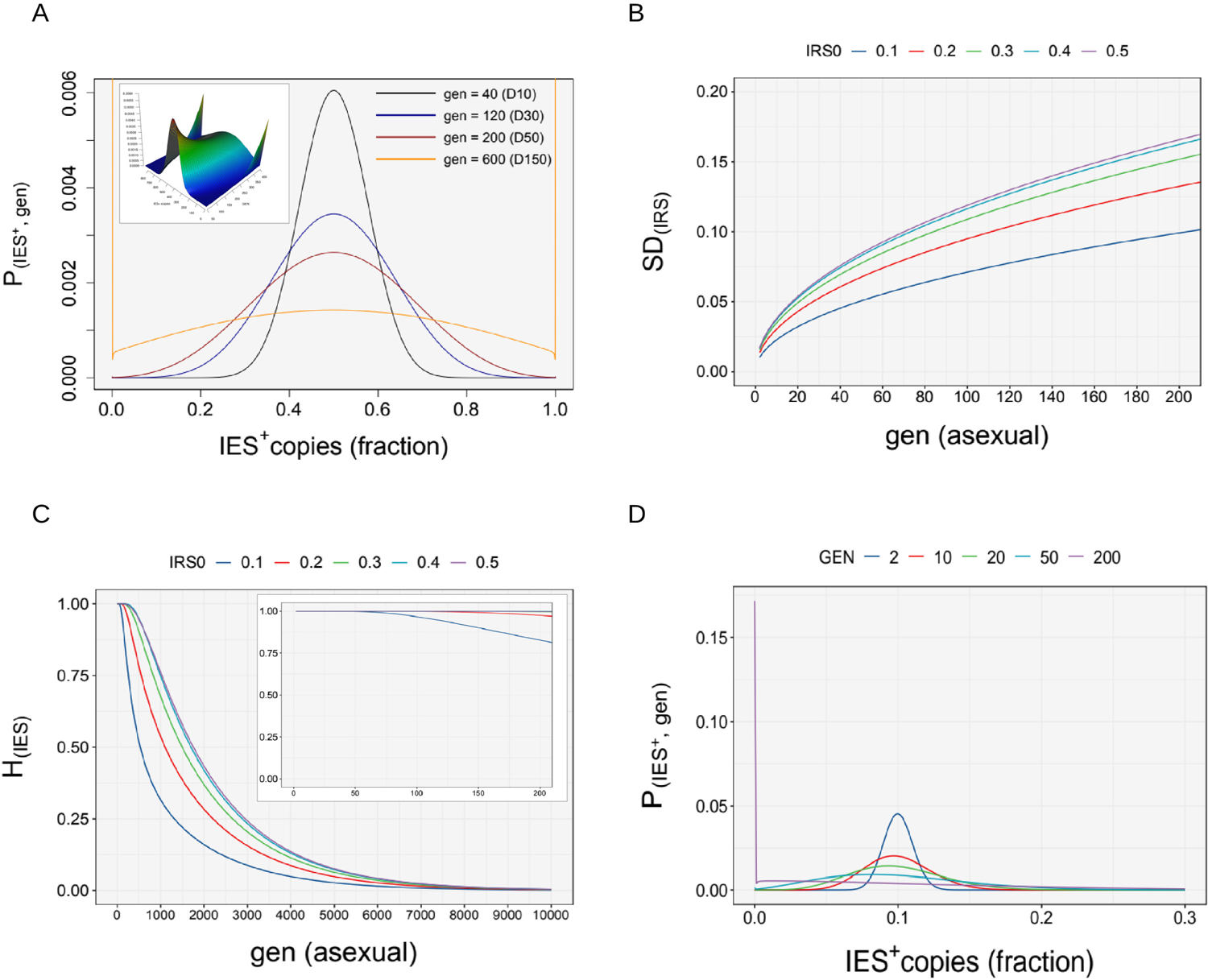
Somatic assortment in P. tetraurelia. A) Probability distribution of IES^+^ copies, P(IES^+^; GEN). Simulated probability distribution of the number of IES^+^ copies in the somatic nuclei after successive amitotic divisions (GEN = 40, 120, 200, 600). Cultivation days (D) are indicated in brackets. The number of IES^+^ copies is expressed as a fraction of the ploidy (k = 860). Simulation is shown for IRS_0_ = 0.5. The inset shows the probability surface across generations. **B)** Effect of assortment on standard deviation, σ(IRS_0_; GEN). Variability of the number of IES^+^ copies due to somatic assortment. The rate of somatic assortment (dσ/dt) is the fastest at IRS_0_ = 0.5, and decreases symmetrically around this value. **C)** Loss of heterozygosity, *H*. Probability of a locus to be in the heterozygous state across divisions. The inset shows the loss of *H* for an IES locus across a full clonal cycle of *P. tetraurelia* (lifespan of ∼200 divisions). **D)** Probability distribution of IES^+^ copies, P(IES^+^; GEN) across amitotic divisions (GEN = 2, 10, 20, 50, 200). Simulation is shown for IRS_0_ = 0.1. For all plots calculations are according to the haploid model. IRS_0_, starting retention levels. GEN, asexual generations. In b and c simulated values are identical for IRS_0_ = [0.1 | 0.9; 0.2 | 0.8; 0.3 | 0.7; 0.4 | 0.6].

Could somatic assortment give rise to phenotypic assortment in *Paramecium*? To address this question, we calculated the fraction of heterozygous cells that after 200 generations would undergo a “phenotypic switch” due to somatic assortment of IESs (*e.g.* IES-bearing gene with IRS0 = 0.5). Assuming an incomplete dominance scenario, wherein gene inactivation occurs when the fraction of IES^+^ copies exceeds 0.85 of the ploidy, only ∼1.4% of the cells (∼6.4% for the chromosomal model) would express the phenotype after 200 divisions (cumulative fraction of cells with IRS >= 0.85 after 200 generations). This fraction becomes smaller when we consider a larger number of assorting somatic chromosomes. It should be emphasized, that the computations reported above refer to single loci. The probability of observing phenotypic assortment increases when considering multiple heterozygous loci simultaneously (roughly estimated by 1-(1-p)^n, n=number of loci, **(Preer 1976)**).

The results of our simulations are consistent with previous indications that somatic assortment in *P. tetraurelia* proceeds rather slowly **(Preer 1976)**. As a consequence, phenotypic assortment is unlikely to be observed within a single clonal cycle **(Nyberg 1976; Preer 1976)**, unless cells exhibit high levels of heterozygosity, which are not characteristic of this self-fertilizing species **(Nanney 1980)** with low nucleotide diversity **(Catania et al. 2009; Johri et al. 2017)**.

### Somatic assortment in *Paramecium*: comparing theoretical and empirical observations

Finally, we compared the experimental dispersion of IES retention levels measured empirically ten (D10) and fourteen (D14) days post self-fertilization with that predicted *in silico*. For the simulations we adopted two models of macronuclear architecture, the *haploid* and the *chromosomal* model, which predict slightly different rates of somatic assortment (see Methods for details and **Figure 3**). We find that on Day 14, the experimental IRS values for a “track set” of highly covered IES loci (n=75; x3 replicates) are slightly more variable than expected, regardless of the model adopted (**Figure 5A** and **Figure 5B**). Namely, 87% (195/225) and ∼90% (202/225) of the empirical IRS values fall within the 95% confidence interval (CI95) predicted by the haploid and chromosomal model, respectively. We find a similar discrepancy between observed and predicted values on Day 10 (**Additional File 1: Figure S4A and Figure S4B**).

**Figure 5.**
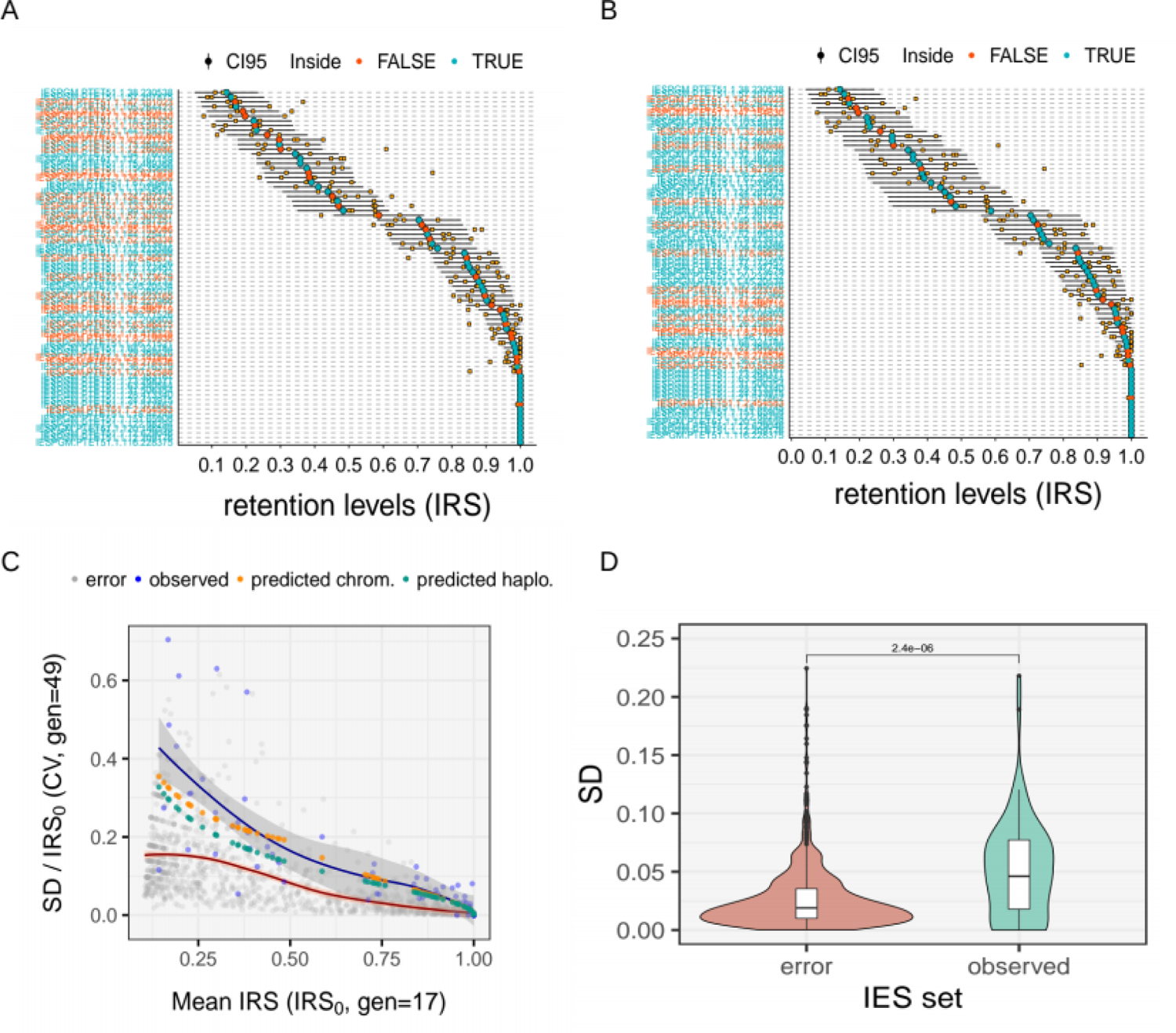
Comparison of observed and theoretical variation of IES retention levels ∼50 amitotic divisions post self-fertilization. A) Haploid model. The empirical distribution of IES retention levels is compared to the theoretical distribution predicted by the haploid model (random assortment of haploid whole-genome subunits). B) Chromosomal model. The empirical distribution of IES retention levels is compared to the theoretical distribution predicted by the chromosomal model (random assortment of chromosomes). Retention levels (orange filled-squares) were measured experimentally with scDNA sequencing 14 days post autogamy (D14, *n*=3) for a selected set of highly covered (> 20 reads) somatic loci (track set, *n*=75). Horizontal black bars represents the theoretical 95% Confidence Interval (CI) constructed on the mean retention levels (IRS_0_, large red or green filled-circles) measured 5 days post autogamy (D5, *n*=4), ∼31 asexual generations prior. Filled-circles (IRS0) are colored in green when the experimentally determined retention level lies inside the 95% CI for all three replicates and red otherwise. IRS, IES Retention Score. C) Observed relative variation of IRSs 14 days post self-fertilization. For each IES, the coefficient of variation of the IRSs measured on day 14 (SDIRS / IRS_0_, gen = 49) is plotted against the mean IRSs measured on day 5 (IRS0, gen = 17). *N* = 75. Predicted IRSs are shown in yellow and green for the *chromosomal* and the *haploid* model simulation, respectively. The distribution of IRS errors (as in Additional File 1: Figure S5) is shown for reference (gray circles). Local polynomial regression is shown in red and blue for the error and the empirical distribution, respectively. D) Comparison of measurement errors with observed IRSs. The absolute random error (SDbIRS) on IRS estimates (*N* = 1,196) is compared to the observed variability of IRSs (SDIRS) measured 14 days post self-fertilization (gen = 49, *N* = 75). Distributions were compared with a Wilcoxon rank sum test. Pairwise comparisons and *P* value is shown above the plot.

We further investigated the relationship between the relative dispersion of retention levels (coefficient of variation, CV) and the starting retention levels (IRS_0_), both for our empirical IRS measurements and the simulated values (**Figure 5C**). The simulations predict a progressive reduction of the coefficient of variation of IES retention levels (CV = SD / IRS0) with increasing starting retention levels (IRS0) (**Figure 5C**). We find that the empirical retention levels measured experimentally 14 days post self-fertilization follow the same pattern, consistent with random assortment of alleles (**Figure 5C**). The observed variability in the empirical IES retention levels could result entirely from the experimental error of the IRS measurements. To test this hypothesis, we quantified the random error of the empirical estimates of IES retention levels (see Methods). We find that although the relative error of the IES retention levels measured experimentally shows a similar reduction with increasing retention levels, this alone cannot explain the observed variation in the empirical IRS estimates (**Figure 5C, Figure 5D and Additional File 1: Figure S5**). More specifically, the observed IRS variation measured 14 days post self-fertilization (gen = 49, *N* = 75) is significantly greater than that from the random error (Wilcoxon rank sum test, *P* = 2.4e-06, effect size r = 0.132 (small), **Figure 5D**). This is consistent with a biological variation of IES retention levels across asexual divisions (as opposed to an experimental artifact).

## Discussion

The study of asexual reproduction in ciliates can provide valuable insights into the evolutionary significance of amitosis. For one, the differentiation of genetically identical heterozygous cells during cell division provides a source of phenotypic plasticity that could facilitate environmental adaptation. For another, selection on somatic assortment could reduce the burden of deleterious germline mutations by preferentially expanding wild type alleles at the expense of the mutated variants **(Zufall et al. 2006)**. As suggested recently, by increasing the fitness variance (boosting selection) in large populations and at the same time dampening the drift load in small populations (Muller’s ratchet), amitosis may even confer “the benefits of sex in the absence of sex” **(Zhang et al. 2019)**.

Here, we studied amitosis and somatic assortment in *Paramecium*, a ciliate that houses ∼860 genome copies in its somatic nucleus **(Allen and Gibson 1972; Woodard et al. 1961)**. Following DNA replication, chromatin sub-units in ciliates are assumed to segregate randomly during amitosis **(Bell 2009; Nanney and Preparata 1979; Orias and Flacks 1975; Preer 1976)**. This implies that the nuclear frequency of an allele in heterozygous clones will change over successive asexual divisions due to stochastic segregation, ultimately resulting in the production of homozygous lines with different phenotypes (*phenotypic assortment*). While there is unequivocal evidence of *phenotypic assortment* in *Tetrahymena* **(Doerder et al. 1992; Merriam and Bruns 1988; Nanney and Preparata 1979; Orias and Flacks 1975)**, the existence of this phenomenon in *Paramecium* is doubtful. In fact, previous experimental evidence argues against its occurrence. In one example, by means of repeated macronucler regeneration in heterozygous clones of *P. aurelia,* Sonneborn was unable to produce phenotypic assortment and suggested that the segregating sub-units be in fact diploid (which is incompatible with assortment) **(Nyberg 1976; Preer 1976; Sonneborn 1947)**. In another, Nyberg used a copper resistance gene as quantitative trait in *P. tetraurelia* and failed to produce evidence for assortment of copper tolerance throughout ∼250 divisions **(Nyberg 1976)**, consistent with Sonneborn’s findings. However, Preer and Nyberg cautioned that higher ploidy levels (>>860n) would still be compatible with random segregation of individual somatic chromosomes **(Nyberg 1976; Preer 1976).** Re-examining the impact that amitosis may have on the somatic variability of *Paramecium* is relevant and particularly timely as it is now clear that potentially heritable somatic variability in *Paramecium* can spark from a fully homozygous state as a consequence of incomplete excision of germline DNA sequences **(Hagen et al. 2020; Vitali et al. 2019)**.

We first explored the extent to which Multiple Displacement Amplification (MDA) coupled with DNA sequencing (which we refer to as scDNA-seq) can be used to faithfully represent the genome of single *Paramecium* cells. To this end, we leveraged whole genome sequencing data from mass culture (bulk DNA-seq) and single *Paramecium* cells obtained from the same clone. We then used scDNA-seq to investigate somatic assortment. We find that scDNA-seq of *Paramecium* AT-rich genomes is affected by mild to moderate positive GC bias (**Figure 1A** and **Table 1, left**). We also uncover a severe representation drop-off near chromosome ends (**Figure 1B** and **Table 1, right**), consistent with the inefficient amplification of template termini in MDA reactions catalyzed by the φ29 DNA polymerase **(Lage et al. 2003; Sabina and Leamon 2015)**. This terminal representation bias could be leveraged to determine the reproducible fragmentation patterns of ciliates’ chromosomes, and/or complement information from telomeric repeats to confirm full-length chromosomes in genome assemblies. In this context, the preferential amplification of large DNA templates in MDA reactions was successfully exploited to preferentially amplify the germline genome of ciliates with highly fragmented somatic DNA **(Maurer-Alcalá et al. 2018)**. Finally, we show that these genome representation biases may result in the underestimation of the number of somatic IESs (due to IES dropout). However, this effect can be ameliorated by increasing sequencing depth (**Table 2, Figure 2 and Additional File 1: Table S1**).

Taking the caveats of scDNA-seq into account, we next assessed the feasibility of tracking somatic assortment of mutant (IES^+^) and wild type (IES*^-^*) alleles across ∼50 asexual generations in single *Paramecium* cells. We tested the degree to which IES retention levels of a “track set” of 75 highly covered loci diverge after 17 and 31 amitotic divisions due to somatic assortment. Our experimental estimates are consistent with a progressive, albeit slow, drift in the fraction of IES^+^ alleles in the nuclei (**Figure 5**). The moderate impact on allele segregation after ∼50 asexual divisions post-fertilization suggests that IESs retention levels are largely sculpted during Programmed DNA Elimination and that amitosis is unlikely to significantly affect allele frequency within a single clonal cycle, at least under the tested conditions, where the power of drift is maximized. Our empirical findings overlap with theoretical expectations based on previously proposed somatic assortment models (**Figure 3**), which we revisit, reproduce (**Additional File 1: Table S3**), and update (**Figure 4**).

Although our empirical observations are compatible with the random segregation of individual chromosome fragments during amitosis (**Figure 5**), at least part of the observed variability of the empirical IES retention levels could result from sources other than somatic assortment, including the measurement errors of retention levels (**Figure 5C** and **Figure S5**) and the progressive fragmentation of somatic chromosomes during clonal senescence **(Gilley and Blackburn 1994)**. Thus, conclusive evidence for the occurrence of somatic assortment in *Paramecium* awaits further experimentation. We anticipate that future experiments to investigate allele segregation in amitotically dividing cells will greatly benefit from the use of scDNA-seq.

In conclusion, we show that single-cell whole-genome sequencing can be successfully used to gain insights into the evolution and structure of AT-rich genomes, provided that the inherent amplification biases of multiple displacement amplification are accounted for. Our study provides a powerful new approach to directly and accurately trace allele segregation in polyploid cells.

## Materials and Methods

### Experiment outline

A single cell of *Paramecium tetraurelia* strain d12 derived from self-fertilization was expanded to a 5 ml mass culture and used as clonal parental population to set up the experiment. The somatic genome of the parental population was purified and sequenced from mass culture (bulk DNA-seq) seven days post self-fertilization (D7) and used as reference. To conduct the single-cell DNA-seq (time-course) experiment, four lines were derived from the parental clonal population and cultured in daily re-isolation regime **(Beisson et al. 2010)**. Single cells were collected in quadruplicates during vegetative growth at five (D5), ten (D10) and fourteen (D14) days post autogamy, and a total of 13 samples (12 scDNA-seq + 1 Bulk DNA-seq) were subjected to Whole Genome Amplification and sequencing. One single cell sample was excluded due to low coverage. Before expanding to mass culture, the progeny of a single sister cell derived from self-fertilization was cultured in daily re-isolation, thus the parental mass culture and the single-cell lines had identical germline genomes and somatic genome configurations before the experiment. Post-autogamous cells of *Paramecium tetraurelia* strain d12 were propagated in isolation cultures at 25 °C as described in **(Vitali et al. 2019)**.

### Amplification biases of MDA-based Whole Genome Amplification

The degree and direction of GC bias from DNA-seq data was evaluated as follows. SAM files were converted to binary, sorted and indexed with SAMtools (version 1.4.1) **(Li et al. 2009)**. Detailed GC bias metrics were collected from mapped reads using the CollectGcBiasMetrics tool of the Picard suite (http://broadinstitute.github.io/picard/). GC bias estimates were calculated as the slope of the linear regression of normalized coverage on GC content between 9 and 50% GC (the two extreme GC content values of *P. tetraurelia*’s genome). For convenience, GC bias estimates are expressed as change of normalized coverage every 10% change in GC content. For a sequencing experiment with mean coverage of 100x, a GC bias of +0.20 corresponds to an increase in coverage of 20 reads every 10% increase in GC content.

The underrepresentation of scaffold ends (here dubbed *terminal bias*) was evaluated as follows. The 115 telomere-capped scaffolds (full-length macronuclear chromosomes) reported in **(Aury et al. 2006)** were selected for the terminal bias analysis. Coverage information was extracted from mapped reads using bedtools (https://bedtools.readthedocs.io/en/latest/index.html). The median base coverage of 49 2kb-overlapping windows (1kb overlap) spanning 50kb from either end of the 115 telomere-capped chromosomes was calculated for each sample. Terminal bias estimates were calculated as the slope of the linear regression (which approximates the true parabolic relationship) of normalized windows coverage on distance from scaffold ends (up to 30kb away from the termini where the increase in coverage plateaus). For convenience, terminal bias estimates are expressed as change of normalized window coverage every 10kb change in distance from chromosome termini. A terminal bias of +0.30 corresponds to an increase in coverage of 30 reads every 10kb increase in distance from the chromosome ends for a sequencing experiment with 100x median base coverage. A FASTQ file was artificially generated from *P. tetraurelia*’s reference genome with ArtificialFastqGenerator **(Frampton and Houlston 2012)** and included as a bias-free reference (aDNA). Multiple samples were processed using custom bash scripts. All data analyses were performed in R **(R Core Team 2020)**.

### IES detection and estimation of retention levels

The extent to which somatic mutations can be detected in the AT-rich genome of *P. tetraurelia* using (MDA-based-) scDNA-seq was evaluated by tracking Internal Eliminated Sequences (IESs) across multiple asexual divisions. IES detection and quantification of their retention levels were performed as in **(Vitali et al. 2019)** using ParTIES **(Denby Wilkes, Arnaiz, and Sperling 2016)**.

### Quantification of the measurement error for IRS estimates

The *random error* of the empirical estimates of IES retention scores (IRSs) was quantified as follows. Briefly, IRSs were estimated genome-wide on all 11 scDNA-seq samples using ParTIES’ MIRET module ran with the *Boundaries* method. For each IES, the module estimates the retention scores on both IES-flanking boundaries (left and right). Low coverage IESs (SUPPORT_MAC + SUPPORT_LEFT + SUPPORT_RIGHT < 20 reads) and IESs with IRSs < 0.1 (IRS_left & IRS_right < 0.1) were removed from the set before downstream analyses. Significant differences between left and right retention levels were tested with a binomial test. *P* values were corrected for multiple testing using the Benjamini–Hochberg procedure. IESs with significantly different left and right retention levels (*Padj* < 0.05, 30 in total) were removed from subsequent analyses to exclude rare events of differential usage of IES boundaries **(Arnaiz et al. 2012; Duret et al. 2008)**. IESs with no variability between left and right scores (304 in total) were also discarded as they represent short IESs whose boundaries are spanned by the same reads (which results in identical scores). A final set of 1,196 IESs was used to estimate the distribution of *random errors* on empirical retention levels. For each IES, the *relative random error* of the retention level was taken as the coefficient of variation of the boundary scores (SD_bIRS_ / bIRS, where bIRS is the mean boundary IRS score).

### Quantification of IES dropout

*Total IES dropout* was calculated as the fraction of all known IES loci (*n*=44,928) with read coverage equal or lower than 20, as a minimum of 20 reads is desirable for robust estimation of IES retention levels (IRS) across most of the IRS spectrum. *Terminal IES dropout* was calculated as the fraction of all known IES loci located within 30 kb from either scaffold ends (*n*=9,986) and with a read coverage equal or lower than 20. A *residual IES dropout*, likely unrelated to amplification biases, is found in the mcDNA sample (see **Table 2**). This term is assumed to scale negatively with the number of read pairs mapped. For any given scDNA sample, the *residual IES dropout* was calculated as the r*esidual IES dropout* found in the mcDNA sample scaled on the sc / mc ratio of mapped read pairs:

*Residual dropout*_sc_ = *Residual dropout*_mc_ / mapped read pairs (sc / mc)

Last, IES dropout attributed to the positive GC bias was calculated as the dropout unexplained by either of the *terminal* or *residual dropout* terms:

*GC dropout* = *Total dropout −* (*Terminal* + *Residual*)

### DNA isolation and sequencing

The somatic genome of the parental population used as a reference to assess the genome representation biases of the scDNA-seq technology was obtained as follows. Somatic nuclei were isolated from a caryonidal mass culture seven days post self-fertilization. ∼10 μg of genomic DNA were purified from 500 ml mass culture in early stationary phase (5*10^5 *Paramecium* cells). The culture was cleaned up by filtration through 8 layers of gauze, cells concentrated on a Nitex filter (Nylon-Netzfilter, 10 μm pore size, 47 mm, Merck KGaA) and pelleted by centrifugation at 800 xRCF for 3 min. Collected cells were stored 1h in Volvic® water before cell lysis to reduce bacterial load. Cells were homogenized in 4 ml of lysis buffer (0.25 M sucrose; 10 mM MgCl2; 10 mM Tris pH 6.8; 0.2% NP40) **(Arnaiz et al. 2012)** by repeated crushing in a syringe barrel (20 ml, 60×25 hypodermic needle). Cell content was washed in 10ml of lysis buffer and macronuclei (MACs) isolated by centrifugation at 1000 xRCF for 15 min at 4°C. Isolated MACs were pre-lysed and gDNA extracted with the NucleoSpin® Tissue Kit following manufacturer’s instructions for DNA isolation from cultured cells.

Single, daily re-isolated *Paramecium* cells strain d12 were washed three times in Volvic® water before DNA amplification and sequencing. Washed cells were subjected to whole genome, Multiple Displacement Amplification (MDA) using the REPLI-g Single Cell Kit (© QIAGEN). The parental somatic DNA from mass culture and whole genome amplification products from single cells (scDNA) were subjected to Paired-End Illumina sequencing (150 bp) on a Novaseq 6000 platform at the Functional Genomic Center Zurich. A total of 12 scDNA samples and 1 bulk DNA sample were sequenced.

### Mathematical Modeling of Somatic Assortment

To model the probability distribution of mutant alleles (IES^+^ copies) across amitotic divisions, we leveraged previously published mathematical models of somatic assortment for ciliates **(Bell 2009; Preer 1976).**

For the *haploid subunit model* we made the following assumptions:

- a.i The ploidy of the somatic nucleus, *k*, is assumed to be 860 **(Allen and Gibson 1972; Woodard et al. 1961)**.
- a.ii The total number of segregating units in the nucleus, *N*, is conserved, and amounts to 2 * *k* (1720) after DNA replication.
- a.iii Each daughter cell receives an equal number of copies, *k*, at each cell division.
- a.iv The number of successes is a natural number ranging from 0 to *k*.
- a.v The process operates in a selection-free environment.

For the *chromosomal model,* we introduced the following modifications:

- a.i We assumed 115 somatic chromosomes (*Chr*)
- a.ii The total number of segregating units, *N,* is conserved and amounts to 2 * *k* * *Chr* (197,800) after DNA replication.
- a.iii Each daughter cell receives an equal number of copies, *N*/*2*, at each cell division.
- a.iv The number of successes is a natural number ranging from 0 to *N*/2.

The following treatment refers to the *haploid model* notation but may be extended to the *chromosomal model* when the modifications reported above are introduced. After a first asexual generation (gen=1), the probability distribution P(*X*) of the number of IES^+^ copies (mutated alleles) per nucleus in the daughter cells (number of successes *x*), represented by the random variable *X*, is a function of the number of IES^+^ copies in the parental nucleus, y_0_, and the number of copies inherited (drawn) upon division, *k*. The number of IES^+^ copies in the parental nucleus (successful elements *m*) available before division (after DNA doubling) equals 2*y*_0_.

P(*X*) is given by the probability mass function of the hypergeometric distribution:

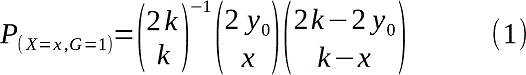

For the following generation, G = t+1, for each *x*, P(*X*, *t*+1) is the summation between *y* = *x*/2 and *y* = (*k*+*x*) / 2 of the product of the probability calculated in (1), denoted P(*X* = *y*, *t*) at G = *t*+1, and the probability of receiving *x* IES^+^ copies, for the range of possible parental IES^+^ copies *y* (*x*/2; (*k*+2)/2) from which *x* could have been arisen. Thus, P(*X*, *t*+1) becomes:

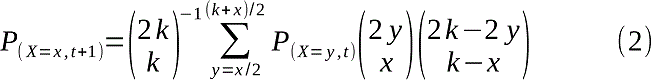

For any given number of successes *x* (number of IES^+^ copies received), the number of IES^+^ copies in the parental nuclei after DNA replication, 2*y*, must have been at least *x*, as the number of IES^+^ copies inherited (*x*) is at most equal to the total number of IES^+^ copies available in the nucleus at the time of division (*x*max = 2*y*), and could have not exceeded *k* + *x*, as *x* is at least equal to the number of IES^+^ copies present in excess with respect to *k*, the number of elements inherited upon division (*x*min = 2*y − k*). Note that the theoretical equivalent of the IES Retention Score (IRS calculated experimentally) is given by IRS = *x*/*k*.

### Rate of somatic assortment

We define the rate of somatic assortment as the change in the standard deviation, σ, of the probability distribution of the fraction of mutated alleles (IES^+^) in the nuclei across sexual generations. At generation t, σ is given by:

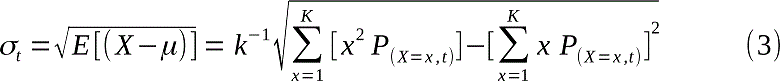

Within 200 divisions (full clonal cycle of *P. tetraurelia*), σ(IRS_0_, t) is approximated by:

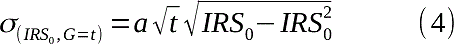

Where IRS_0_ (the starting parental retention level) can assume values between 0.1 and 0.9 in steps of 0.1, and the parameter *a* is equal to 0.0245 (1.4201*0.0245 for the chromosomal model). Thus, for each IRS_0_, the (instantaneous) rate of somatic assortment is the derivative function of σ with respect to t calculated as follows:

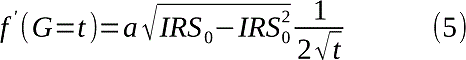

### Rate of loss of Heterozygosity

The process of somatic assortment eventually leads to the complete loss of the heterozygous state, with nuclei containing only either mutated (IES^+^) or wild type alleles (IES*^-^*). The rate of loss of heterozygosity due to somatic assortment is calculated as the change of the cumulative probability of the heterozygous state, *H*, across asexual generations. *H*, at generation *t*, is given by:

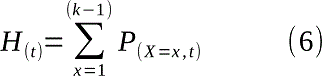

Both the haploid and chromosomal models assume that the total number of segregating units is conserved and that each of the two daughter cells receives exactly half that amount at each division. However, in the chromosomal model the total number of copies of a given locus is not fixed and the *number* of IES^+^ copies will slowly tend toward a third absorbing boundary (in addition to only IES^+^ or only IES^-^ copies): no copies of either alleles (*nullisomic* locus). Nevertheless, as this tendency toward chromosomal loss will affect both alleles, we assumed the relative *fraction* of IES^+^ copies (retention level) to remain symmetrical. Equation (4) is a previously-unpublished mathematical equation determined through evolutionary searches performed with the A.I.-powered modeling engine Eureqa (https://www.nutonian.com/products/eureqa/).

### Bioinformatic simulation of somatic assortment

Through bioinformatic simulations we estimated the probability distribution of mutated alleles (P(*X*)), its standard deviation (σ), and the fraction of heterozygous cells (*H)*, across successive asexual generations. We simulated the process for the daily re-isolation and mass culture regimes, with daily bottlenecks of 1 and 2^12^ (4096) cells (for a culture of ∼50 ml), respectively. The assumptions to model somatic assortment were identical to those made for the mathematical simulation with the *haploid model* (a.i – a.v). The 860 binary subunits (two parental haplotypes) were represented with binary digits (bits). The simulation was started with an input ratio of 0.5 (430 zeros and 430 ones). Cell division was simulated by drawing an equal number of subunits (860 bits) without replacement from a single set (G2 cell, 1720 bits), followed by partitioning into two sets (daughter cells). For each iteration (day) of simulated isolation culture (daily re-isolation), a single, randomly selected founder cell was used to start a series of 4 successive *in silico* cell divisions (4 div. / Day), which produced 2^4^ (16) cells. The process was repeated 2^10^ (1,024) times to simulate replicate isolation cultures, for a total of 2^14^ (16,384) cells (N = 2^4^ * 2^10^ = 2^14^) across replicates. In contrast, for each iteration (day) of simulated mass culture, 2^10^ (1,024), randomly selected founder cells (*inocolum*) were used to commence a series of 4 successive *in silico* cell divisions, which produced a total of 2^14^ (16,384) cells (N = 2^10^ * 2^4^ = 2^14^). The simulation was protracted for 200 generations.

### Experimental estimates of somatic assortment

To study somatic assortment experimentally, we sequenced the somatic genome of single cells using scDNA-seq across ∼50 asexual divisions (see Experiment outline). Cells divided on average ∼3.5 times per day (25°C) in all single cell lines studied. IRS values were determined experimentally at Day 5 (*gen* ∼17), Day 10 (*gen* ∼35) and Day 14 (*gen* ∼49). To account for the amplification biases introduced by the MDA reaction, a set of somatic IESs (IRS > 0.1 at Day 5) with coverage greater than 20 reads shared by all 11 scDNA samples was selected (track set, *n* = 75) for further analysis.

### Simulation of retention levels and confidence intervals

The mean retention levels measured experimentally 5 days post self-fertilization (D5, gen = 17, *n* = 4) were taken as starting retention levels (IRS_0_) to initiate the somatic assortment simulation. The probability distribution of the fraction of IES copies (simulated IRSs) expected at generation ∼35 (D10) and ∼49 (D14) was calculated individually for each IES locus in the track set (*N* = 75). The predicted standard deviation (σ) was calculated from the simulated probability distribution using equation 3. σ was then used to construct a 95% Confidence Interval (CI95) around IRS_0_ for each of the 75 IES loci in the track set. Due to the high ploidy of *P. tetraurelia* (∼860), the simulated IRS probability distributions approximate the normal distribution within the ∼50 asexual generations investigated in this study (for 0.1 < IRS_0_ < 0.9). Thus, the CI95 was calculated for Day 10 (Day 10 − Day 5, ∼17 *gen*), and Day 14 (D14 − D5, ∼31 *gen*) as IRS_0_ ± 2*σ(IRS_0_, *gen*) (0 ≤ x ≤ 1), with σ being a function of IRS_0_ and the number of generations occurred (under the adopted model, equation (3-4)).

## Data availability

All DNA reads generated in this study are available in the European Nucleotide Archive (https://www.ebi.ac.uk/ena/browser/home) under the study accession number PRJEB43365. All data generated or analyzed during this study will be provided as Supplementary Information files.

## Code availability

All custom R scripts associated with this submission will be provided as Supplementary Code. The software used to simulate the random segregation of genetic elements in polyploid nuclei **(Vitali et al. 2021)** is available at https://doi.org/10.5281/zenodo.4573521 licensed under the MIT license.

## Acknowledgment

This work was carried out within the research training group ‘Evolutionary Processes in Adaptation and Disease funded by the Deutsche Forschungsgemeinschaft (DFG, German Science Foundation) - 281125614/GRK 2220. We wish to thank Andrea Vitali who suggested the A.I.-powered modeling engine Eureqa for equation discovery.

## Contributions

**V.V.** performed the experiments; wrote the manuscript; analyzed the data; performed mathematical and bioinformatic simulations; developed software and designed the experiments. **R.H.** performed exploratory data analyses; processed DNA samples. **F.C.** conceived and designed the project; supervised; wrote the manuscript; and secured funding.

## Competing Interests

The Authors declare no competing interests.

## Notes

https://doi.org/10.5281/zenodo.4573521

https://github.com/biowalter/Amitosis

### Competing Interest Statement

The authors have declared no competing interest.

