## Additional File 1 for "Fifty generations of amitosis: tracing asymmetric allele segregation in polyploid cells with single-cell DNA sequencing"

### --- Additional File 1: Supplementary Figures and Tables ---

**Running title**: Investigating amitosis via single-cell DNA sequencing

**Keywords:** Amitosis, single-cell DNA sequencing, developmental variation, copy number variation, somatic mutations, somatic assortment, polyploidy

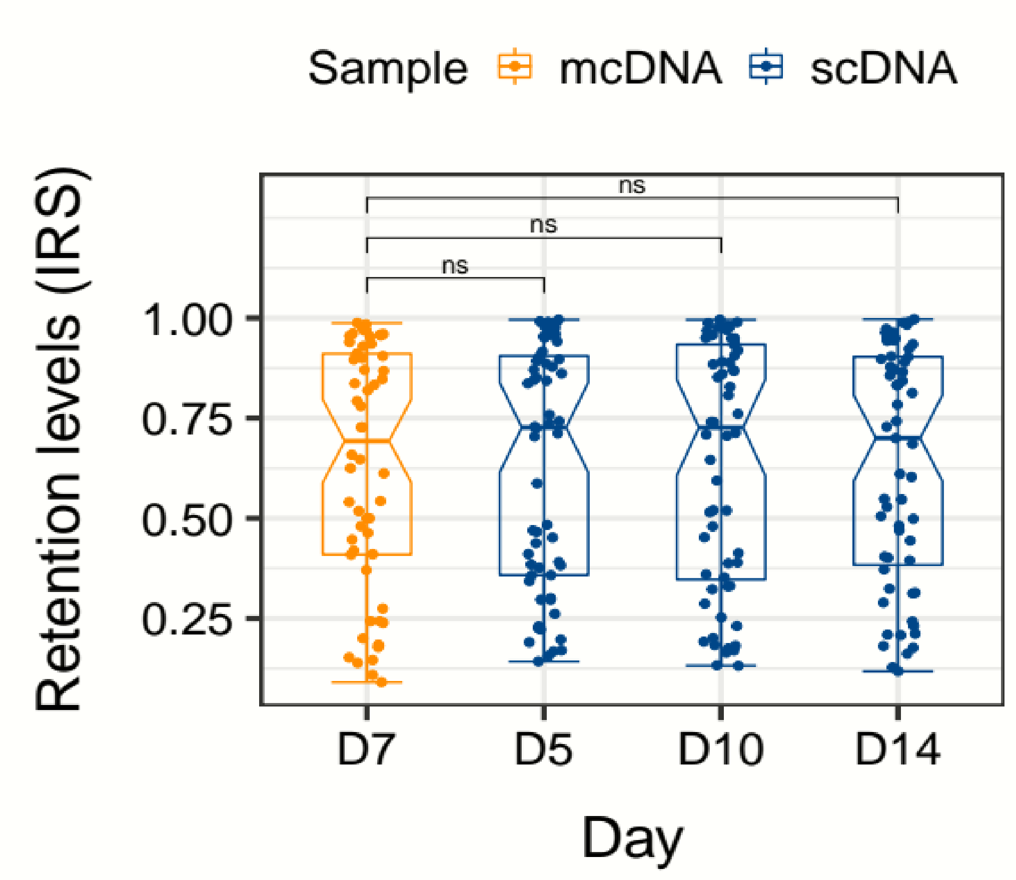


**Figure S1. Comparison of empirical IES retention levels between bulk DNA-seq from mass culture and DNA-seq from single cells.** A set of highly covered (>20 reads) somatic IESs shared across scDNA samples (*n*=75, "Track Set") was selected for comparison. scDNA samples were collected in quadruplicates on each day (D5, D10, D14). IES retention levels (IRS) for scDNA samples were averaged out across replicates. IESs with IRS = 1 were excluded. IES sample size after filtering: D7 (*n*=58); D5 (*n*=60); D10 (*n*=60); D14 (n=59). IRS distributions were compared with a Wilcoxon rank sum test. mcDNA was used as reference for comparison. Significance levels for pairwise comparisons are shown above each plot (ns: *Padj* > 0.05). IRS, IES retention score / retention level. IES, Internal Eliminated Sequences. Somatic IESs, IESs from macronuclear DNA with IRS >= 0.1. mcDNA, bulk DNA-seq from mass culture. scDNA, DNA-seq from single-cell whole-genome amplification products (MDA reactions). MDA, Multiple, Displacement Amplification.

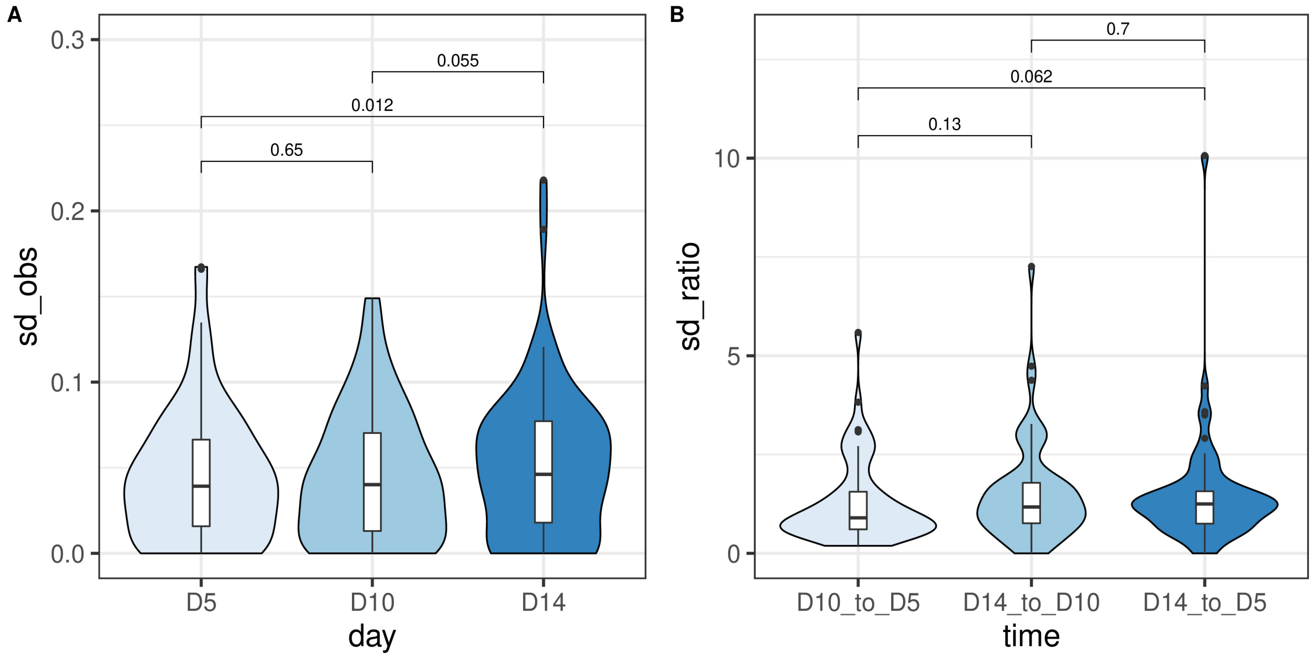


**Figure S2. Variability of retention levels across time. A)** Standard deviation of empirical retention levels (observed $SD_{IRS}$) 5, 10 and 14 days after self-fertilization (D5, D10, D14) for a highly supported set of IESs (*N* = 75). $SD_{IRS}$ values were computed across replicate scDNA samples (D5, *n* = 4, D10, *n* = 4, D14, *n* = 3). **B)** Standard deviation ratios for empirical retention levels (observed $SD_{IRS}$) for a highly supported set of IESs. SD ratios ($SDR_{IRS}$) were computed pairwise between time points (D14 to D5, D14 to D10, D10 to D5). *N* = 60. Distributions were compared with a Wilcoxon signed rank test. Pairwise comparisons and *P* are shown above each plot.

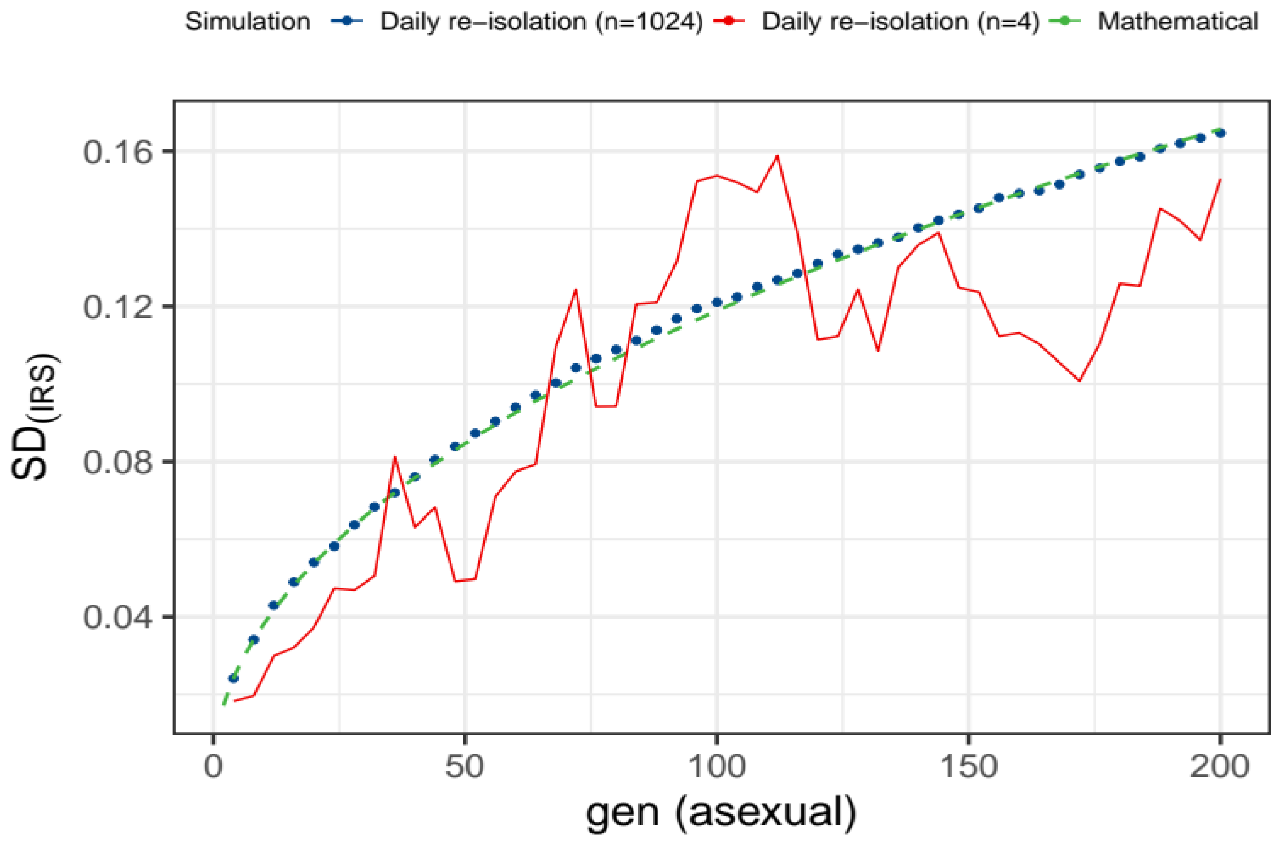


**Figure S3. Validation of mathematical modeling through bioinformatic simulation of somatic assortment.** Change in standard deviation (SD) of IES retention levels (IRS) across asexual generations as predicted by bioinformatic and mathematical simulations of somatic assortment. All predictions are based on the *haploid model*. See Methods for details on the simulations.

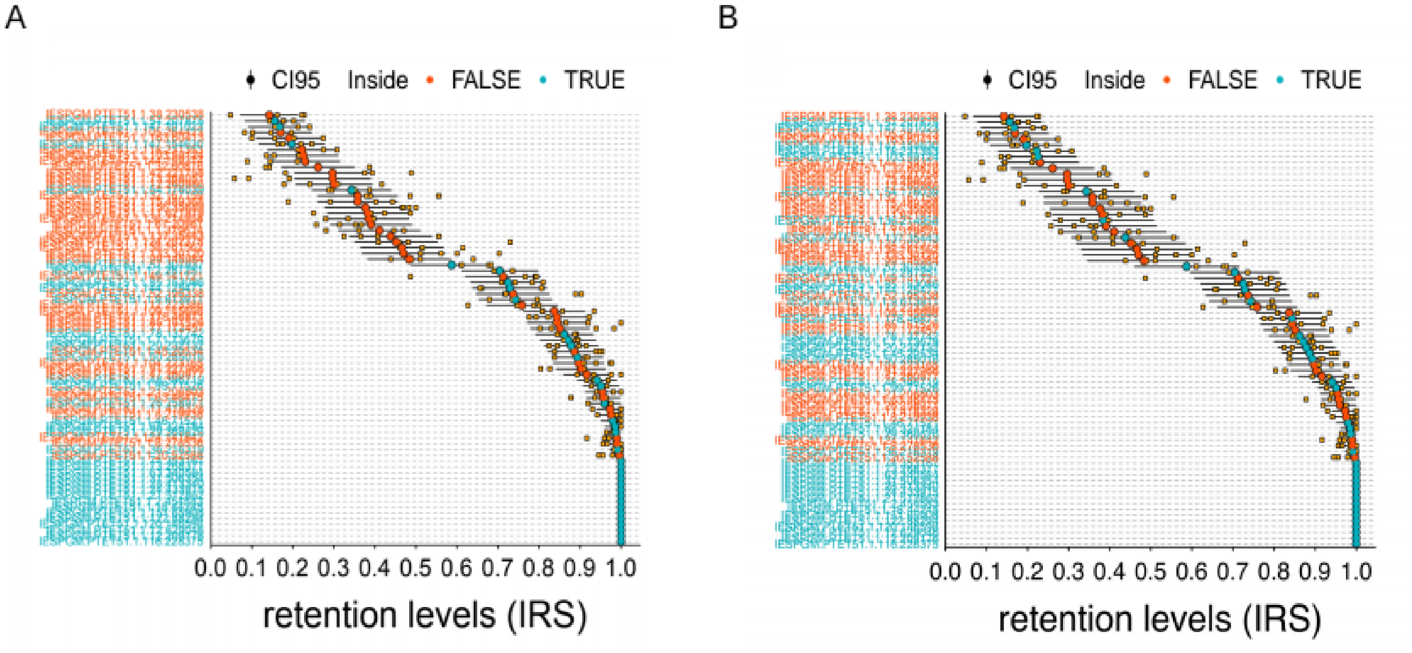


**Figure S4. Observed and theoretical variation of IES retention levels after ~35 amitotic divisions. A) Haploid model.** The empirical distribution of IES retention levels is compared to the theoretical distribution predicted by the haploid model (random assortment of haploid whole-genome subunits). **B) Chromosomal model.** The empirical distribution of IES retention levels is compared to the theoretical distribution predicted by the chromosomal model (random assortment of chromosomes).

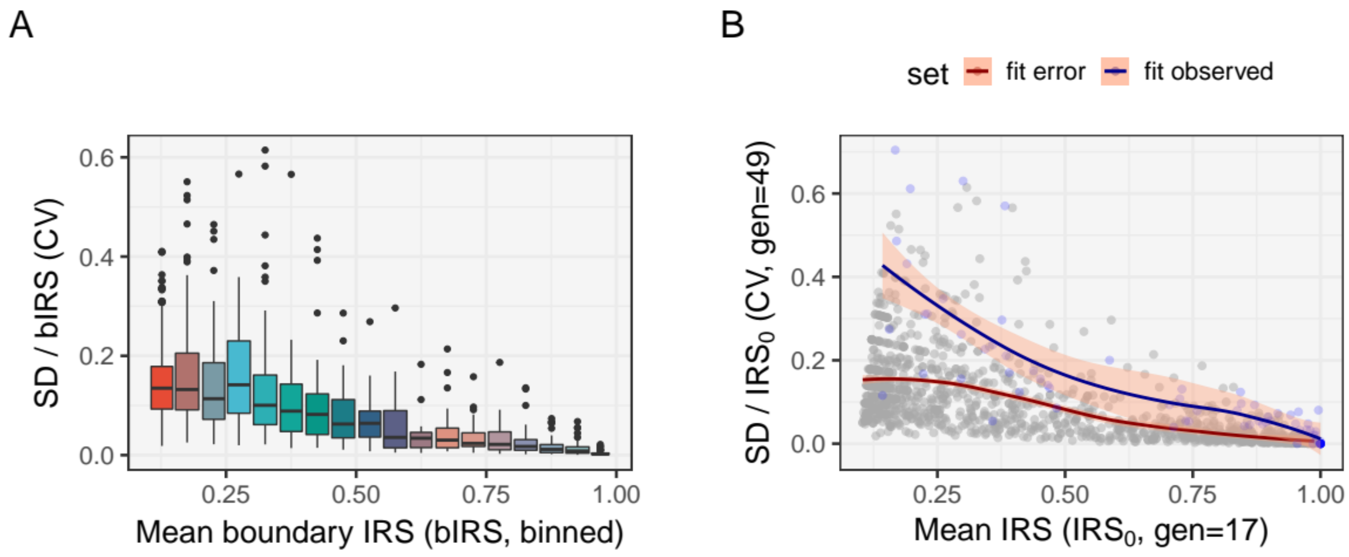


**Figure S5. Random error distribution for IRS measurements. A)** Relative error of IRS measurements (binned, size = 0.05) across IRS values. For each IES, the coefficient of variation of the boundary scores ($SD_{bIRS}$ / bIRS) is plotted against the mean boundary score (bIRS). *N* = 1,196 (11 scDNA samples). $SD_{bIRS}$ values were computed across left and right boundary scores (see Methods for details). Summary statistics for the relative random error distribution are: 1st Qu. = 0.0196562, Median = 0.0656980, Mean = 0.0937853, 3rd Qu. = 0.1355395. **B)** Observed relative variation of IRSs 14 days post self-fertilization (blue circles). For each IES, the coefficient of variation of the IRSs measured on day 14 ($SD_{IRS}$ / $IRS_0$, gen = 49) is plotted against the mean IRSs measured on day 5 ($IRS_0$, gen = 17). *N* = 75 (D5, *n* = 4; D14, *n* = 3). The distribution of IRS errors (as in A) is shown for comparison (gray circles, not binned). Local polynomial regression is shown in red and blue lines for the error and the empirical distribution, respectively. Summary statistics for the absolute random error distribution are: 1st Qu. = 0.0099702, Median = 0.0189858, Mean = 0.0266522, 3rd Qu. = 0.0354437.

**Table S1. Genome coverage statistics.** Coverage statistics for computer generated reads (aDNA) mass culture (mcDNA) and single-cell sequencing (scDNA) samples. Total pairs (M), total number of read pairs (millions). Mapped pairs (M), total number of mapped read pairs (millions). Mapping rate, fraction of reads mapped to the reference genome. Coverage, average number of per-base mapped reads. Average scaffold coverage, number of per-base mapped reads average d across scaffolds (Coverage = average scaffold coverage weighted on scaffold size). Coverage 20 (%), proportion of bases in the genome covered with at least 20 reads (expressed in percentage). Average scaffold coverage 20 (%), proportion of bases in a scaffold covered with at least 20 reads averaged out across scaffolds (expressed in percentage). A threshold of 20 reads was taken as reference as accurate estimation of small IES retention levels requires at least 20 reads. aDNA, artificially-generated DNA sequencing. mcDNA, mass culture DNA sequencing. scDNA, single-cell DNA sequencing. scDNA_1x, scDNA samples with approximately the same number of mapped reads compared to the mcDNA sample (5*10^6 < *n*° of mapped reads < 15*10^6, *n*=6). scDNA_2x, scDNA samples with approximately twice as many mapped reads compared to the mcDNA sample (*n*° of mapped reads > 19*10^6, *n*=4).

| **Sample** | **Total pairs (M)** | **Mapped pairs (M)** | **Mapping rate** | **Genome** | | **Scaffolds** | |
|---|---|---|---|---|---|---|---|
| | | | | **Coverage (reads)** | **Coverage 20 (%)** | **Average coverage (reads)** | **Average coverage 20 (%)** |
| **aDNA** | 9.99 | 9.99 | 1.00 | 41.57 | 99.02 | 41.43 | 90.68 |
| **mcDNA** | 13.56 | 10.92 | 0.80 | 45.23 | 95.89 | 31.39 | 53.97 |
| **scDNA 1x** | 12.02 ± 3.47 | 11.29 ± 3.62 | 0.93 ± 0.05 | 46.27 ± 14.69 | 83.57 ± 8.26 | 23.85 ± 8.18 | 34.01 ± 5.87 |
| **scDNA 2x** | 20.77 ± 0.027 | 19.67 ± 0.51 | 0.94 ± 0.024 | 81.00 ± 1.97 | 92.72 ± 1.54 | 39.20 ± 0.089 | 43.86 ± 3.17 |

**Table S2. Empirical and theoretical estimates of IES retention levels across asexual divisions.** Empirical, observed variation in IES retention levels (IRS). Haploid, calculations based on the haploid whole-genome subunits model. Chromosomal, calculations based on the chromosomal model. Median standard deviation of IRS values is shown within brackets. Mean retention levels at Day 5 were taken as starting retention levels ($IRS_0$). Data are relative to 75 highly covered IES loci (> 20 mapped reads). IRS, IES Retention Scores. Day, 5, 10 and 14, ~17, ~35 and ~49 divisions after self-fertilization respectively. $SD_{IRS}$, observed and predicted standard deviation of retention levels.

| | Time Point | | |
|---|---|---|---|
| Mean $SD_{IRS}$ (*median*) | **Day 5** | **Day 10** | **Day 14** |
| **Empirical** | 0.0449 (*0.0392*) | 0.0449 (*0.0401*) | 0.0507 (*0.0461*) |
| **Haploid** | nd | 0.0296 (*0.0356*) | 0.0394 (*0.0474*) |
| **Chromosomal** | nd | 0.0394 (*0.0506*) | 0.0559 (*0.0673*) |

**Table S3. Predictions of somatic assortment-generated variability in allele frequency distribution across 250 divisions according to this study and Preer 1976.** Predictions with the *haploid model* are for an initial number of 430 subunits and a total number of 860. Predictions with the *chromosomal model* are for an initial number of 430 subunits and a total number of 36,980 (860 * 43). The standard deviation of the allele frequency is reported as fraction of the starting number of segregating subunits as in Preer 1976 (rather than fraction of the ploidy level as reported by SENES.py (this study)). The discrepancy between SENES.py and Preer's predictions for the *chromosomal model* is due to our assumption that the tendency toward chromosomal loss will affect both alleles and thus the relative fraction of IES+ copies (retention level) would remain symmetrical. To drop this assumption and reproduce Preer's predictions exactly, SENES.py should be ran with the --nullisomics flag on.

| | *Haploid model* | | *Chromosomal Model* | |
|---|---|---|---|---|
| **GEN** | **SD (SENES.py)[a]** | **SD (Preer 1976)[b]** | **SD (SENES.py)[c]** | **SD (Preer 1976)[d]** |
| 50 | 0.16934 | 0.17000 | 0.23960 | 0.24000 |
| 100 | 0.23776 | 0.24000 | 0.33576 | 0.34000 |
| 150 | 0.28911 | 0.29000 | 0.40283 | 0.42000 |
| 200 | 0.33146 | 0.33000 | 0.45346 | 0.48000 |
| 250 | 0.36795 | 0.37000 | 0.49376 | 0.54000 |

[a] Simulation with -m haploid -k 860 -i 0.5 -g 250

[b] Data from Table 5

[c] Simulation with -m chromosomal -c 43 -k 860 -i 0.5 -g 250

[d] Data from Table 3
